# The novel Coronavirus enigma: Phylogeny and mutation analyses of SARS-CoV-2 viruses circulating in India during early 2020

**DOI:** 10.1101/2020.05.25.114199

**Authors:** Anindita Banerjee, Rakesh Sarkar, Suvrotoa Mitra, Mahadeb Lo, Shanta Dutta, Mamta Chawla-Sarkar

**Author notes:** Corresponding Author: Dr. Mamta Chawla-Sarkar, Division of Virology, ICMR-National Institute of Cholera and Enteric Diseases, P-33, C.I.T. Road, Scheme-XM, Beliaghata, Kolkata-700010, West Bengal, India. These authors contributed equally to this work (Anindita Banerjee, Rakesh Sarkar).

## Abstract

**Background:** This is a comprehensive analysis of 46 Indian SARS-CoV-2 genome sequences available from the NCBI and GISAID repository during early 2020. Evolutionary dynamics, gene-specific phylogeny and emergence of the novel co-evolving mutations in nine structural and non-structural genes among circulating SARS-CoV-2 strains in ten states of India have been assessed.

**Materials and methods:** 46 SARS-CoV-2 nucleotide sequences submitted from India were downloaded from the GISAID (39/46) or from NCBI (7/46) database. Phylogenetic study and analyses of mutation were based on the nine structural and non-structural genes of SARS-CoV-2 strains. Secondary structure of RdRP/NSP12 protein was predicted with respect to the novel A97V mutation.

**Results:** Phylogenetic analyses revealed the evolution of “genome-type clusters” and adaptive selection of “L” type SARS-CoV-2 strains with genetic closeness to the bat SARS-like coronaviruses than pangolin or MERS-CoVs. With regards to the novel co-evolving mutations, 2 groups are seen to circulate in India at present: the “major group” (52.2%) and the “minor group” (30.4%), harboring four and five co-existing mutations, respectively. The “major group” mutations fall in the A2a clade. All the minor group mutations, except 11083G>T (L37F, NSP6) were unique to the Indian isolates.

**Conclusion:** The study highlights rapidly evolving SARS-CoV-2 virus and co-circulation of multiple clades and sub-clades, driving this pandemic worldwide. This comprehensive study is a potential resource for monitoring the novel mutations in the viral genome, changes in viral pathogenesis, for designing vaccines and other therapeutics.

## 1. Introduction

COVID-19 pandemic caused by the novel Coronavirus SARS-CoV-2 initially reported from Wuhan, China in December 2019, spread across the world within 3 months [1]. As of 23^rd^ May 2020, more than 5.3 million have tested positive with death toll of approximately 340,411 deaths in more than 210 countries. Phylogenetic analyses reveal that SARS-CoV-2 clusters within the subgenus Sarbecovirus under the genus Betacoronavirus, and has probably undergone zoonotic transmission from the bats through the possible intermediate host Malayan pangolins, culminating among humans [2]. The positive sense, single-stranded RNA genome of SARS-CoV-2 is continuously mutating resulting in circulation of multiple clades within short span of time (December’2019-May’2020). Thus dissecting the evolutionary characteristics, nature of variations in different strains and understanding the single nucleotide polymorphisms of the SARS CoV-2 virus is the need of the hour. Previous reports on genetic and evolutionary dynamics have tried to deduce transmission dynamics through which virus made its way into human from bats during early phase of pandemic, but many questions remain unanswered even though more sequence data is made available. Studying the heterogeneous genomic constellations within specific geographical settings would help to understand its complex epidemiology and formulate region specific strategies.

The first three cases from India were reported in Kerala during January 2020 having travel history to Wuhan [3]; subsequently 69,597 active cases, 51,783 recovered cases and 3720 deaths have been officially recorded in India as on 23^rd^ May, 2020, 08:00 IST GMT+5:30 [4]. Inspite of high population density, poor hygiene conditions and overburdened health care system, death rate due to SARS-COV-2 is lower in India compared to USA or European countries [5]. Thus to understand the phylodynamics of circulating strains in India, this study was initiated to analyze the complete viral genome sequences submitted in NCBI GenBank and Global Initiative on Sharing All Influenza Data (GISAID) [6], from 46 SARS-CoV-2 strains circulating across 10 differentially affected states within India. In order to elucidate the possible ancestry, gene-wise phylogeny of the Indian strains has been deciphered with respect to other isolates reported from Europe, USA and China along with Coronavirus strains belonging to other genera infecting humans and other animal hosts. The novel co-evolving mutations among the Indian SARS-CoV-2 strains have also been analyzed.

Through this genome-analyses and phylogenetic approach we would throw light on the natural evolution of SARS-CoV-2 from its existing ancestors in zoonotic reservoirs. Furthermore, analyzing the novel mutations accumulated in the viral genome over the period of time with reference to Wuhan strains (clade O) may shed light on their impact on protein structure and function.

## 2. Material and Methods

### 2.1. Sequence mining

46 SARS-CoV-2 nucleotide sequences submitted from India were downloaded either from the GISAID (39/46) or from NCBI (7/46) database for phylogenetic analyses. Several other reference gene sequences of SARS-CoV-2 as well as other types of Coronaviruses were downloaded from NCBI submitted from other countries, for dendrogram construction and further lineage analyses. 7 sequences from Karnataka (India) were excluded for RdRP/NSP12 gene analysis due to low nucleotide sequence coverage.

### 2.2. Phylogenetic analyses and screening of mutations

Nine phylogenetic dendrograms were constructed with respect to 2 structural genes (Spike and Nucleocapsid) and 7 non-structural genes (NSP2, NSP3, NSP4, NSP6, NSP7, NSP8, and NSP12). Multiple sequence alignment for all the respective set of gene sequences was done using MUSCLE v3.8.31. Amino acid sequences were deduced through TRANSEQ (Transeq Nucleotide to Protein Sequence Conversion Tool, EMBL-EBI, Cambridgeshire, UK). Phylogenetic dendrograms were constructed by MEGA, version X (Molecular Evolutionary Genetics Analysis), using the maximum-likelihood statistical method (at 1000 bootstrap replicates), using the best fit nucleotide substitution models for each dendrogram. The best fit models were determined through model testing parameter of MEGAX. Various novel mutations analyzed among the Indian isolates in comparison to the prototype SARS-CoV-2 strain from Wuhan (MN908947.3/SARSCOV-2 Wuhan-Hu-1).

### 2.3. Secondary structure prediction of RdRp having A97V mutation

We used CSFFP (Chou and Fasman Secondary Structure Prediction) online server to predict the secondary structure of RdRP/NSP12 with novel A97V mutation [7].

## 3. Results

### 3.1. Phylogenetic Analysis of the Structural and Non-Structural Genes

#### 3.1.1. Spike gene (S)

The phylogenetic dendrogram for S gene revealed that the 46 Indian study isolates clustered among themselves within the same lineage of betacoronavirus SARS-CoV-2 in eight different sub-clusters (a-h), while one strain from Telangana (EPI ISL 431101) extruded out singly. Sub-cluster a (5 strains), b (6 strains), c (6 strains), d (5 strains), e (5 strains), f (7 strains), g (5 strains), were close to clade specific strains A5, B1, A3, A2, A1a and A2a, respectively; while cluster h (6 strains) was close to A1, B2, B4-1 and B4-2 clade-specific strains (Fig.1A). The sub-cluster b strains grouped with SARS-CoV-2 isolated from a Tiger in New York zoo; while h-strains clustered with another carnivorous mammal Mink SARS-CoV-2. All the representative Indian strains had 99-100% nucleotide sequence homology among themselves. The prototype SARS-CoV-2 strain belonging to the O clade (MN908947.3/SARS-COV-2/HUMAN/CHN/Wuhan-Hu-1/2019) was present in the same lineage with the Indian strains in between sub-cluster g and h (>99% identity). The Indian strains had 92.8-93% and 83.5% homology with Bat (EPI_ISL_402131/COV/BAT/YUNNAN/RATG13/2013) and Pangolin coronavirus (EPI_ISL_410540/COV/PANGOLIN/GUANGXI/ P5L/2017), respectively. Homology was much less (75.8-76.7%) with other bat SARS-like coronavirus strains like MG772933.1/SARS-LIKE-COV/BAT/BAT-SL-COVZC45/2017 and MG772934.1/SARS-LIKE-COV/BAT/BAT-SL-COVZXC21/2015; while MERS-Coronaviruses (KJ713299.1/MERS-COV/CAMEL/SAU/KSA-CAMEL-376/2013 and KU308549.1/MERS-COV/HUMAN/KOR/SEOUL-SNU1-035/2015) were distantly related to the Indian SARS-CoV-2 strains (52.5-52.9% identity).

**Figure 1(A):**
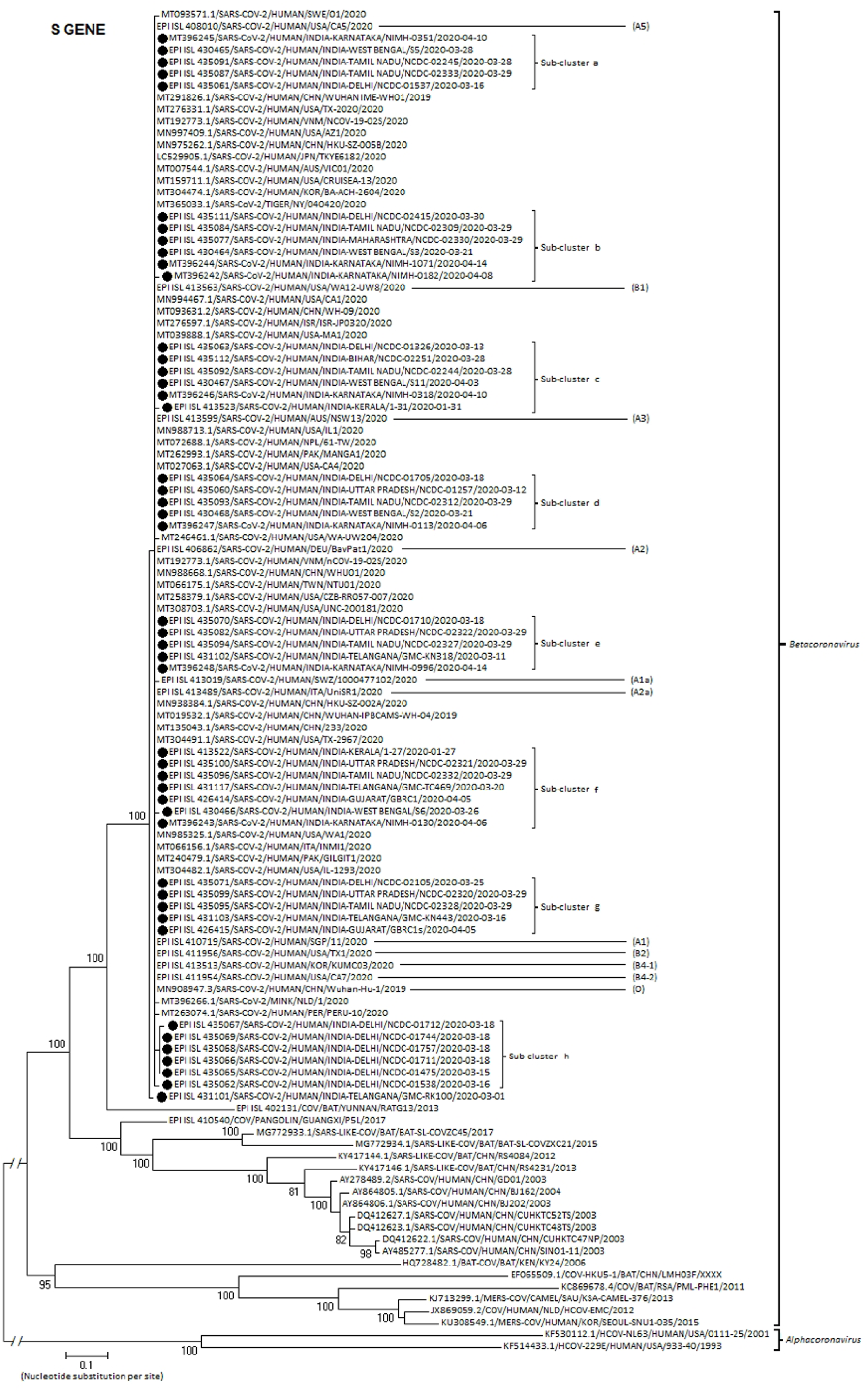
Molecular Phylogenetic analysis by Maximum Likelihood method: Phylogenetic dendrogram based on nucleotide sequences of Spike (S) gene of SARS-CoV-2 strains circulating in India during early 2020, with other known strains of respective genotype. The representative Indian strains have been marked with a solid circle (●). Scale-bar was set at 0.1 nucleotide substitutions per site. Bootstrap values of less than 70% are not shown. The best fit model which was used for constructing the phylogenetic dendrogram was General Time Reversible model with Gamma distribution having Invariant sites (GTR+G+I).

#### 3.1.2. Nucleocapsid gene (N)

The phylogenetic dendrogram for N gene showed clustering of the 46 Indian study isolates among themselves within the same lineage of betacoronavirus SARS-CoV-2 in three different sub-clusters (a, b and c).Three strainsfrom Telangana (EPI ISL 431101), Karnataka (MT396245.1) and Delhi (EPI ISL 435111) extruded out separately, close to sub-cluster b strains. Sub-cluster a (5 strains) were exclusively from Delhi. All the clade-specific strains (A1, A1a, A2, A2a, A3, A5, B2, B1,B4-1 and B4-2) as well as the prototype SARS-CoV-2 strain-O clade (MN908947.3/SARS-COV-2/HUMAN/CHN/Wuhan-Hu-1/2019) clustered near sub-cluster c (9 strains); while the major sub-cluster b (29 strains) were distant to any of the clade-specific strains (Fig.1B). All the representative Indian strains had >99.8% nucleotide identity among themselves as well as with the different clade-specific strains. The Indian strains had 91-97% sequence identity with Bat coronaviruses (EPI_ISL_402131/COV/BAT/YUNNAN/RATG13/2013; MG772933.1/SARS-LIKE-COV/BAT/BAT-SL-COVZC45/2017 and MG772934.1/SARS-LIKE-COV/BAT/BAT-SL-COVZXC21/2015) and 91% similarity with Pangolin strains (EPI_ISL_410540/COV/PANGOLIN/GUANGXI/ P5L/2017); while MERS-Coronaviruses (KJ713299.1/MERS-COV/CAMEL/SAU/KSA-CAMEL-376/2013 and KU308549.1/MERS-COV/HUMAN/KOR/SEOUL-SNU1-035/2015) genetically far-less related to the Indian SARS-CoV-2 strains (56.9-57.3% identity).

**Figure 1(B):**
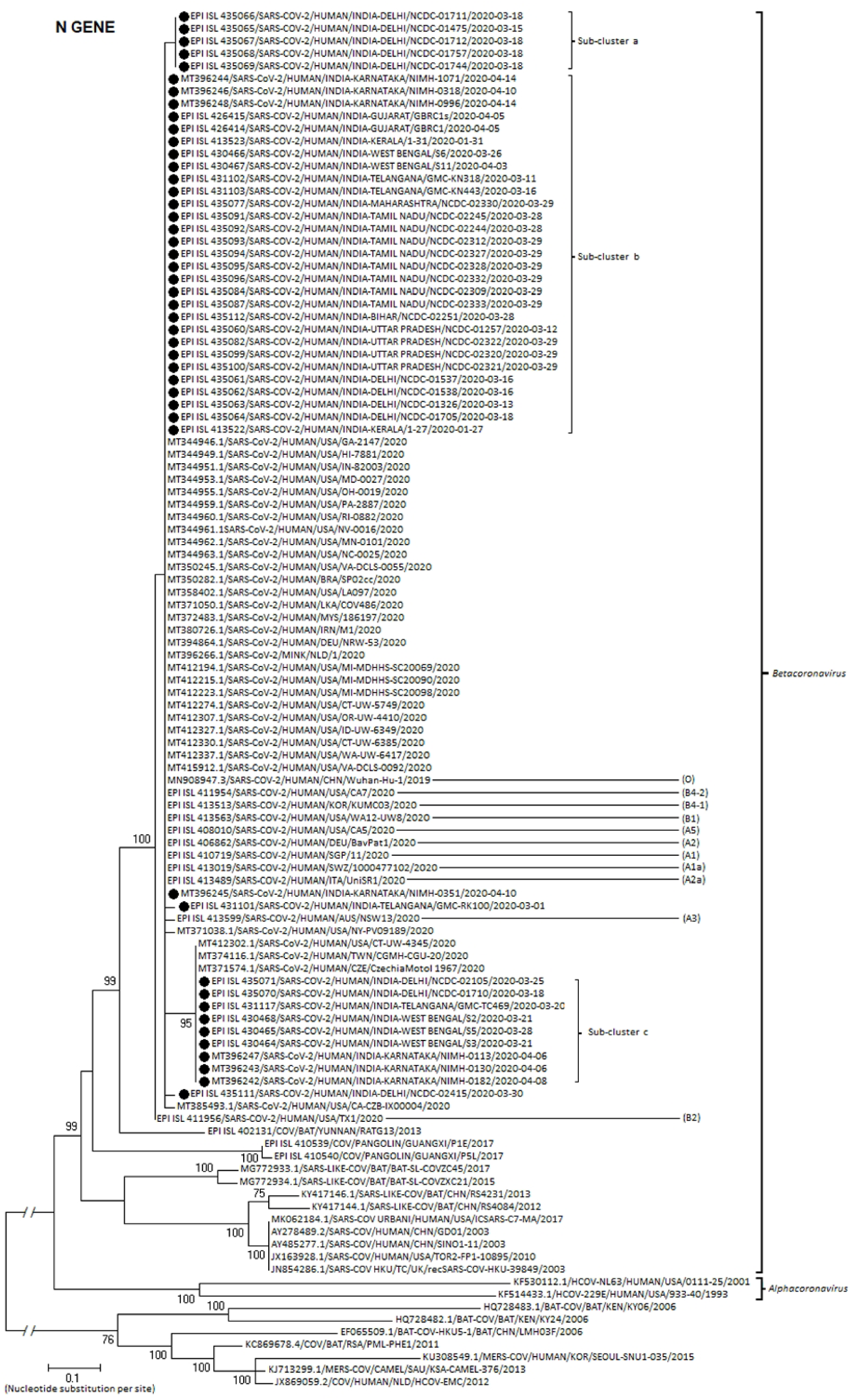
Molecular Phylogenetic analysis by Maximum Likelihood method: Phylogenetic dendrogram based on nucleotide sequences of Nucleocapsid (N) gene of SARS-CoV-2 strains circulating in India during early 2020, with other known strains of respective genotype. The representative Indian strains have been marked with a solid circle (●). Scale-bar was set at 0.1 nucleotide substitutions per site. Bootstrap values of less than 70% are not shown. The best fit model which was used for constructing the phylogenetic dendrogram was General Time Reversible model with Gamma distribution having Invariant sites (GTR+G+I).

#### 3.1.3. RNA-dependent RNA polymerase gene (NSP12)

The phylogenetic dendrogram for NSP12 gene showed clustering of the 39 Indian study isolates among themselves within the same lineage of betacoronavirus SARS-CoV-2 into two sub-clusters (a and b). Four strains, one from Telangana (EPI ISL 431101), other from Delhi (EPI ISL 435111)-close to A1a clade-strain, while two from Kerala (EPI ISL 413522-close to A3 and EPI ISL 413523-close to B1, B2, B4-1 and B4-2 clade-strains) were placed distant to these 2 sub-clusters in the dendrogram. Sub-cluster a (20 strains) and b (15 strains) were close to clade-specific strains A2a and A2, respectively (Fig.1C). All the Indian strains had >99.8% nucleotide identity among themselves as well as the different clade-specific strains. The prototype SARS-CoV-2 strain-O clade (MN908947.3/SARS-COV-2/HUMAN/CHN/Wuhan-Hu-1/2019) was distant to both the sub-clusters. The Indian strains had 97.8% sequence homology with Bat coronavirus (EPI_ISL_402131/COV/BAT/YUNNAN/RATG13/2013); while 86.7-88.6% similarity with both Pangolin (EPI_ISL_410540/COV/PANGOLIN/GUANGXI/ P5L/2017) and other bat SARS-like coronavirus strains (MG772933.1/SARS-LIKE-COV/BAT/BAT-SL-COVZC45/2017 and MG772934.1/SARS-LIKE-COV/BAT/BAT-SL-COVZXC21/2015). MERS-Coronaviruses (KJ713299.1/MERS-COV/CAMEL/SAU/ KSA-CAMEL-376/2013 and KU308549.1/MERS-COV/HUMAN/KOR/SEOUL-SNU1-035/2015) were distantly related to the Indian SARS-CoV-2 strains (68.1% identity).

**Figure 1(C):**
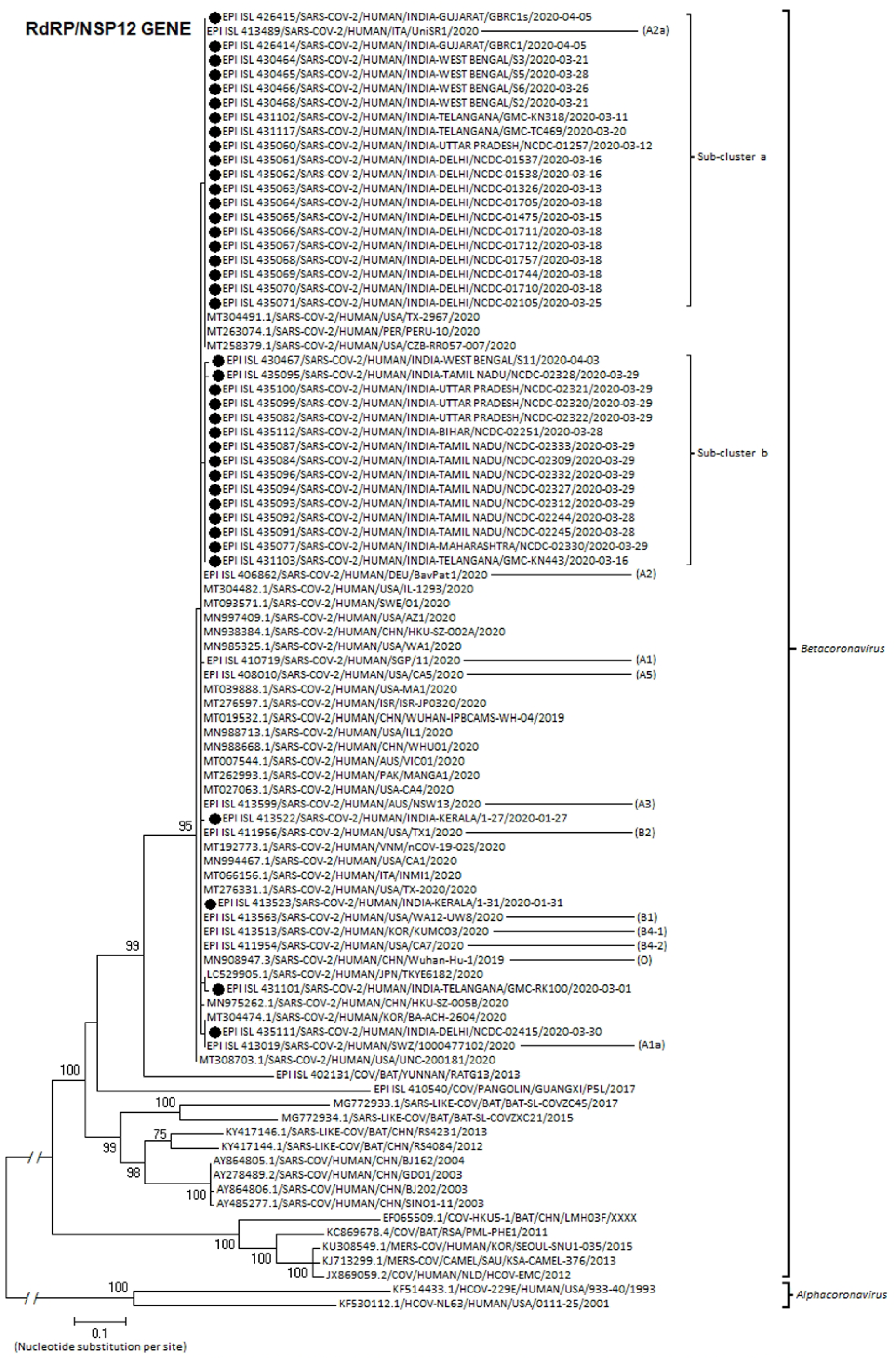
Molecular Phylogenetic analysis by Maximum Likelihood method: Phylogenetic dendrogram based on nucleotide sequences of RNA dependent RNA Polymerase/NSP12 gene of SARS-CoV-2 strains circulating in India during early 2020, with other known strains of respective genotype. The representative Indian strains have been marked with a solid circle (●). Scale-bar was set at 0.1 nucleotide substitutions per site. Bootstrap values of less than 70% are not shown. The best fit model which was used for constructing the phylogenetic dendrogram was General Time Reversible model with Gamma distribution having Invariant sites (GTR+G+I).

#### 3.1.4. NSP2, NSP3, NSP4, NSP6, NSP7 and NSP8 genes

The dendrograms of all these six genes showed a similar pattern. All the 46 Indian strains clustered in a single group among themselves within the betacoronavirus lineage of SARS-CoV-2. All the representative Indian strains had 99.9-100% DNA homology among themselves. The prototype SARS-CoV-2 strain-O clade (MN908947.3/SARS-COV-2/HUMAN/CHN/Wuhan-Hu-1/2019) clustered near the Indian cluster (99.9% identity); while rest of the clade-specific strains (A1, A1a, A2, A2a, A3, A5, B1, B2, B4-1, B4-2) were also present close to the study isolates under the same lineage (>99% homology). Only 1 strain in NSP2 dendrogram (EPI_ISL 435060) extruded out singly close to the prototype strain. SARS-CoV-2 strains isolated from carnivorous mammals like Mink and Tiger also grouped closed to these strains in the dendrogram (99.9% identity) (Fig 1D-1I). The Indian cluster revealed 95.4-98.1% nucleotide sequence similarity with bat coronavirus EPI_ISL_402131/COV/BAT/YUNNAN/RATG13/2013; while Pangolin derived strain EPI_ISL_410540/COV/PANGOLIN/GUANGXI/P5L/2017 showed less identity (83-87.5%). MERS-CoV strains NC_019843.3/MERS-COV/HUMAN/NLD/HCOV-EMC/2012 and KU740200.1/MERS-COV/CAMEL/EGYPT/NRCE-NC163/2014 exhibited a long phylogenetic distance (only 49.6-60.8% homology) to the Indian isolates.

**Figure 1(D):**
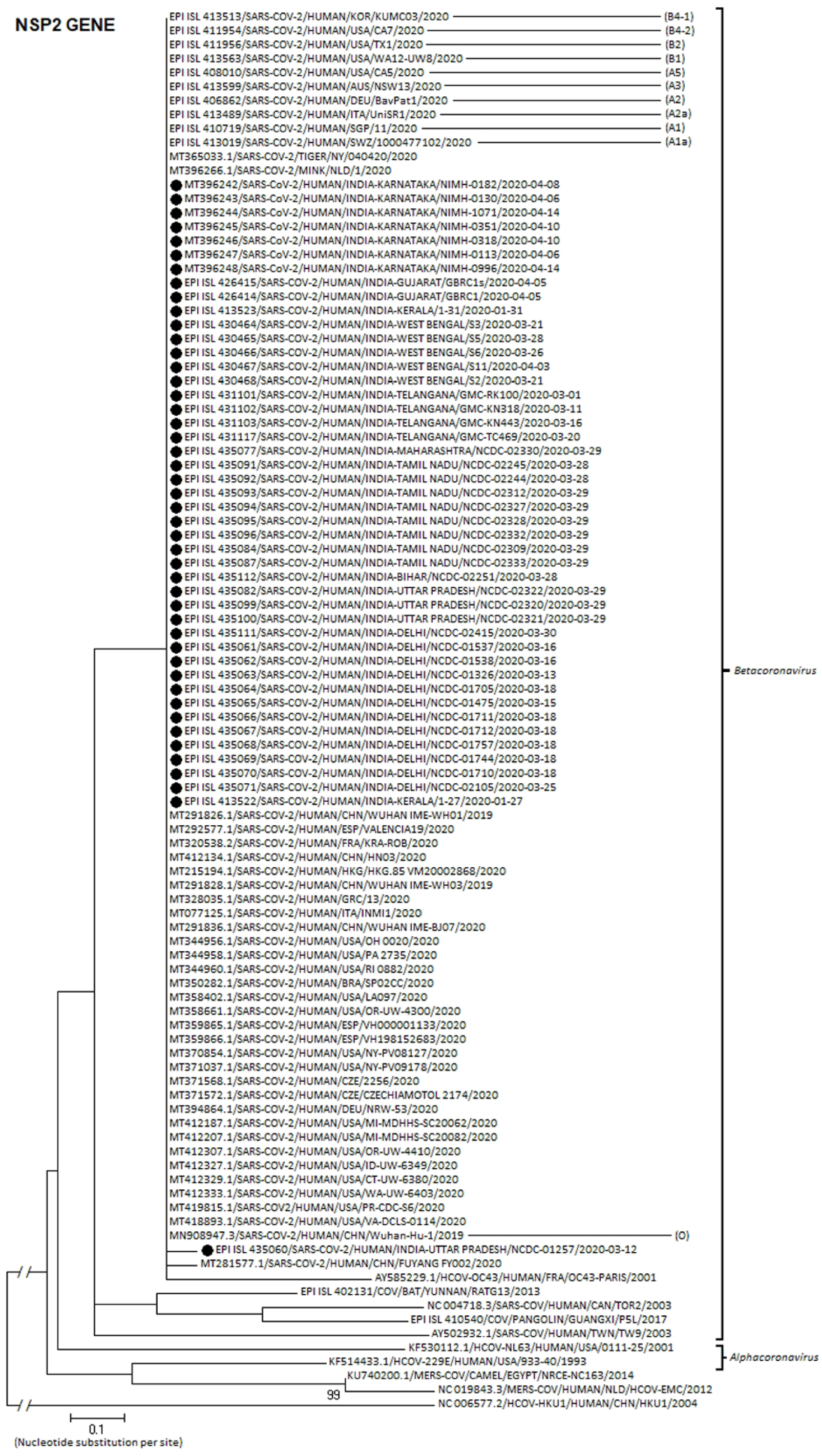
Molecular Phylogenetic analysis by Maximum Likelihood method: Phylogenetic dendrogram based on nucleotide sequences of NSP2 gene of SARSCoV-2 strains circulating in India during early 2020, with other known strains of respective genotype. The representative Indian strains have been marked with a solid circle (●). Scale-bar was set at 0.1 nucleotide substitutions per site. Bootstrap values of less than 70% are not shown. The best fit model which was used for constructing the phylogenetic dendrogram was General Time Reversible model with Gamma distribution having Invariant sites (GTR+G-I).

**Figure 1(E):**
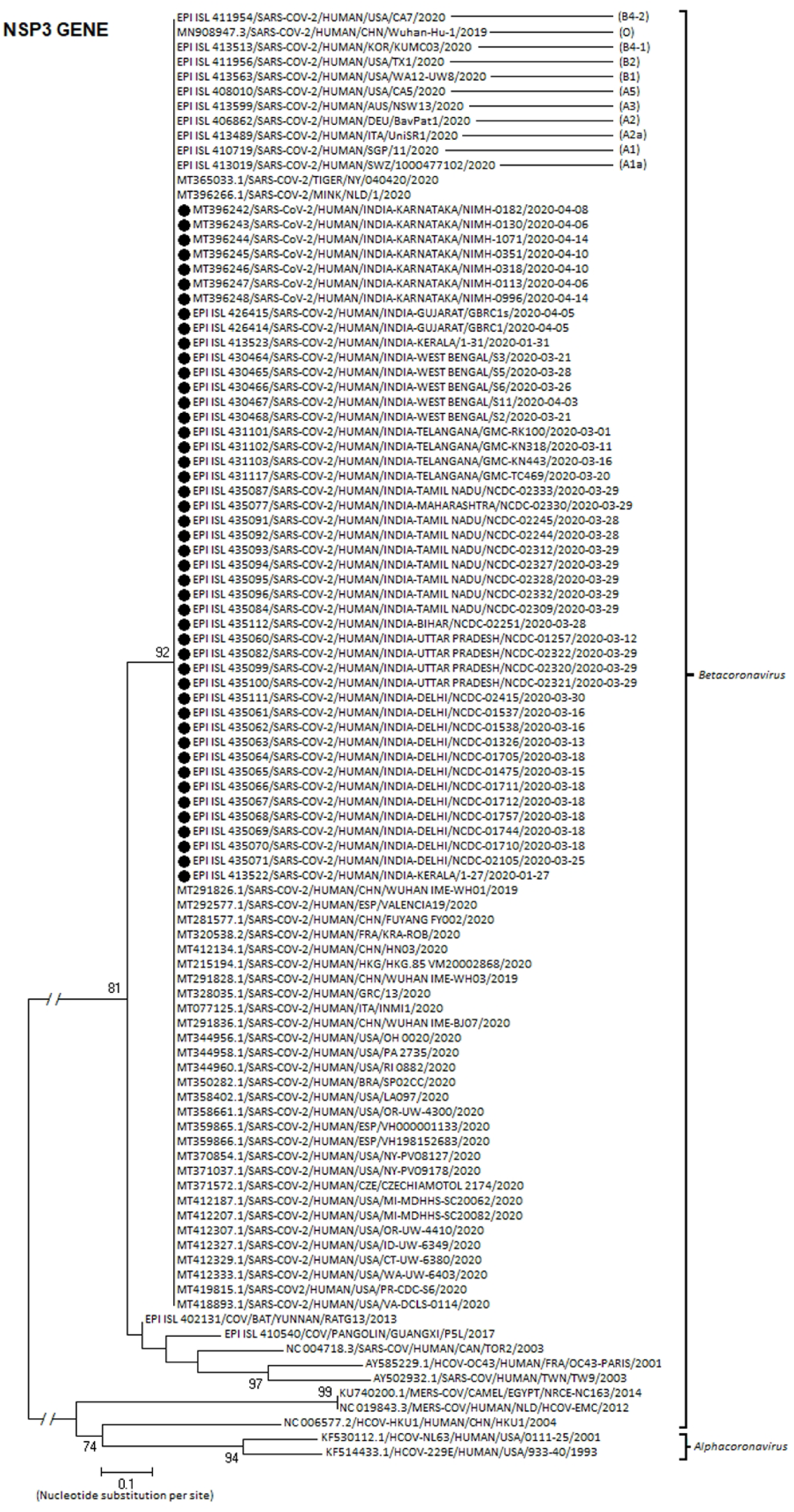
Molecular Phylogenetic analysis by Maximum Likelihood method: Phylogenetic dendrogram based on nucleotide sequences of NSP3 gene of SARS-CoV-2 strains circulating in India during early 2020, with other known strains of respective genotype. The representative Indian strains have been marked with a solid circle (●). Scale-bar was set at 0.1 nucleotide substitutions per site. Bootstrap values of less than 70% are not shown. The best fit model which was used for constructing the phylogenetic dendrogram was Hasegawa-Kishino-Yano model (HKY).

**Figure 1(F):**
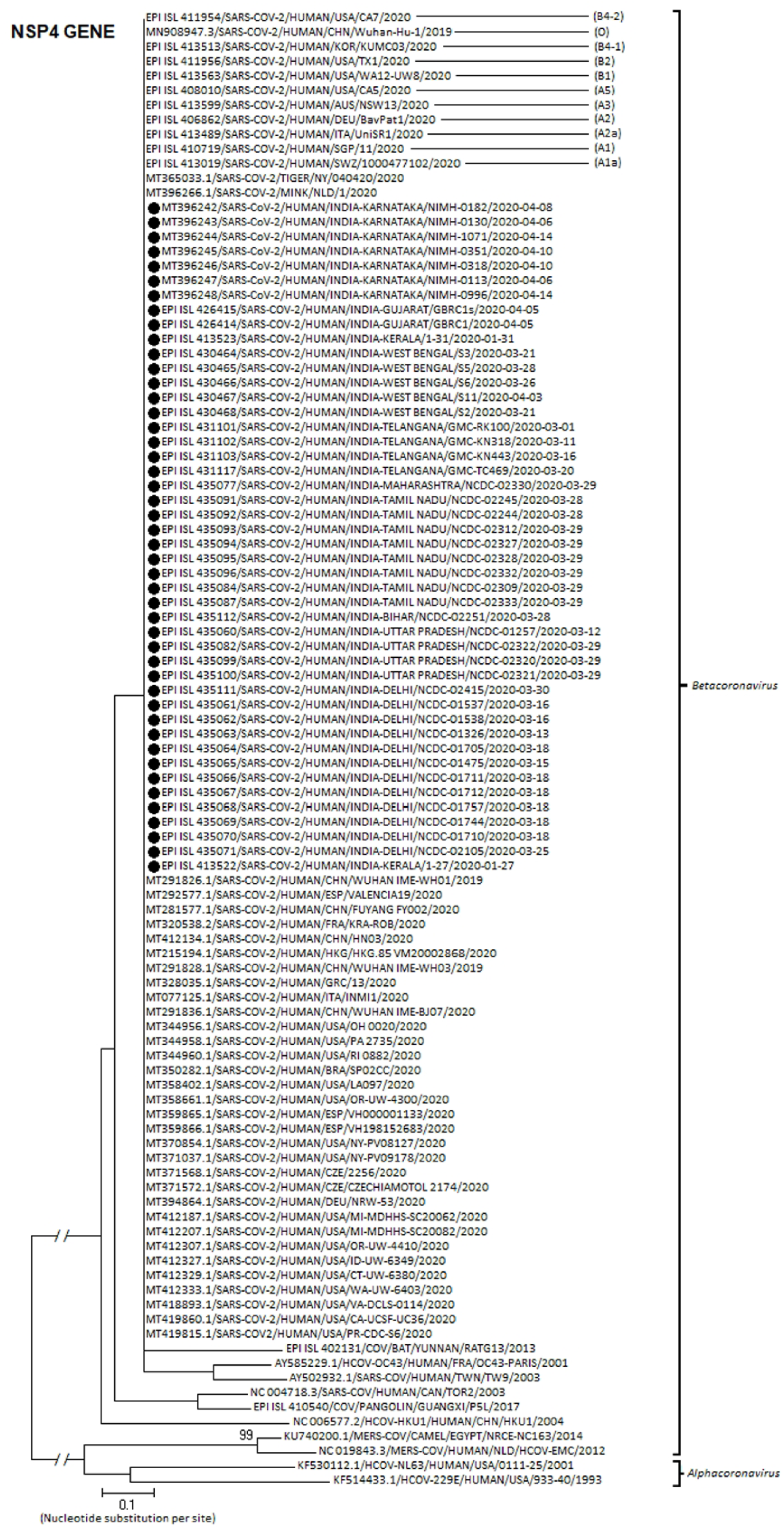
Molecular Phylogenetic analysis by Maximum Likelihood method: Phylogenetic dendrogram based on nucleotide sequences of NSP4 gene of SARS-CoV-2 strains circulating in India during early 2020, with other known strains of respective genotype. The representative Indian strains have been marked with a solid circle (●). Scale-bar was set at 0.1 nucleotide substitutions per site. Bootstrap values of less than 70% are not shown. The best fit model which was used for constructing the phylogenetic dendrogram was Tamura-Nei (TN93) model.

**Figure 1(G):**
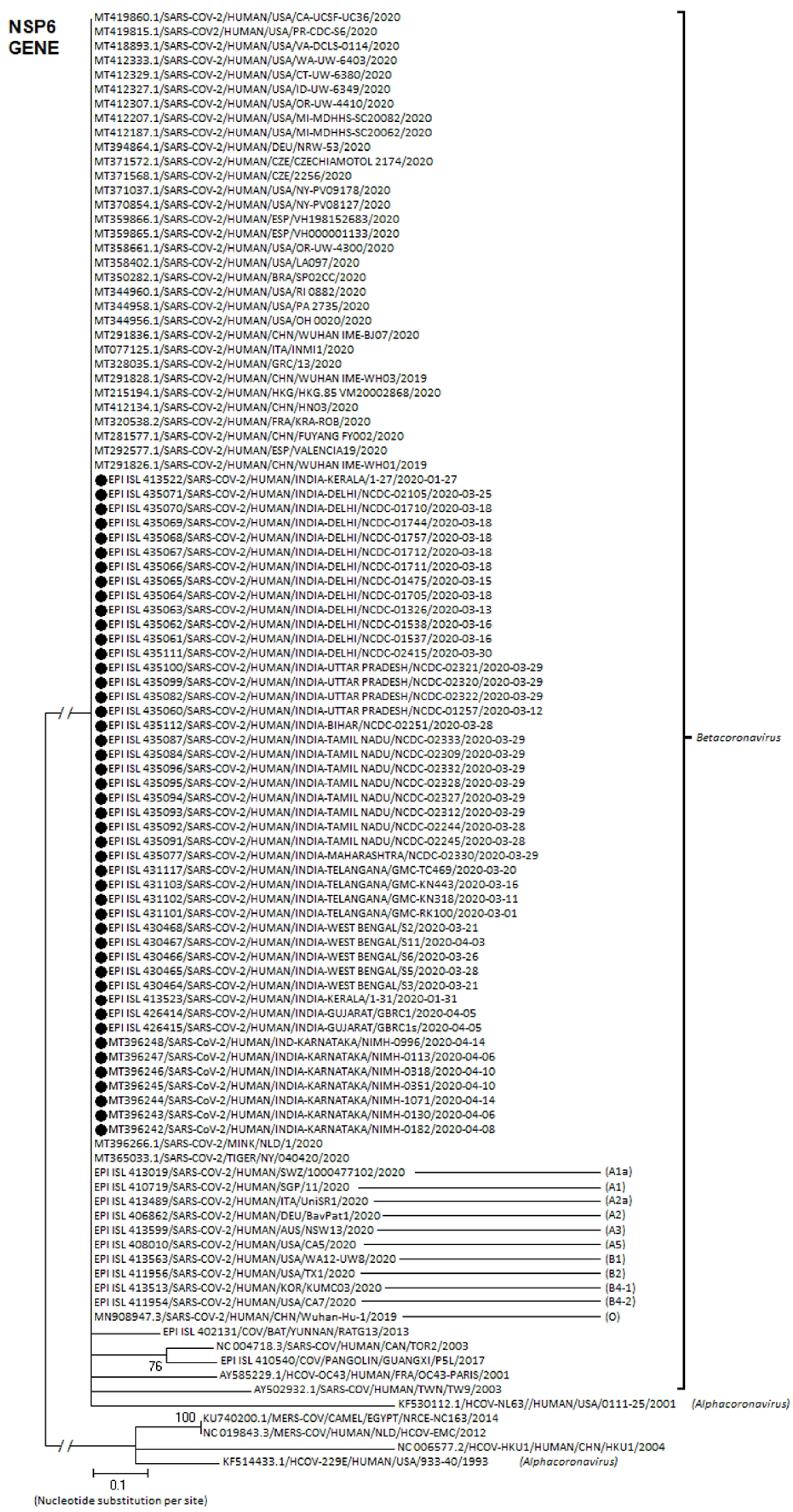
Molecular Phylogenetic analysis by Maximum Likelihood method: Phylogenetic dendrogram based on nucleotide sequences of NSP6 gene of SARS-CoV-2 strains circulating in India during early 2020, with other known strains of respective genotype. The representative Indian strains have been marked with a solid circle (●). Scale-bar was set at 0.1 nucleotide substitutions per site. Bootstrap values of less than 70% are not shown. The best fit model which was used for constructing the phylogenetic dendrogram was Tamura-3 model with Gamma distribution having Invariant sites (T92+G+I).

**Figure 1(H):**
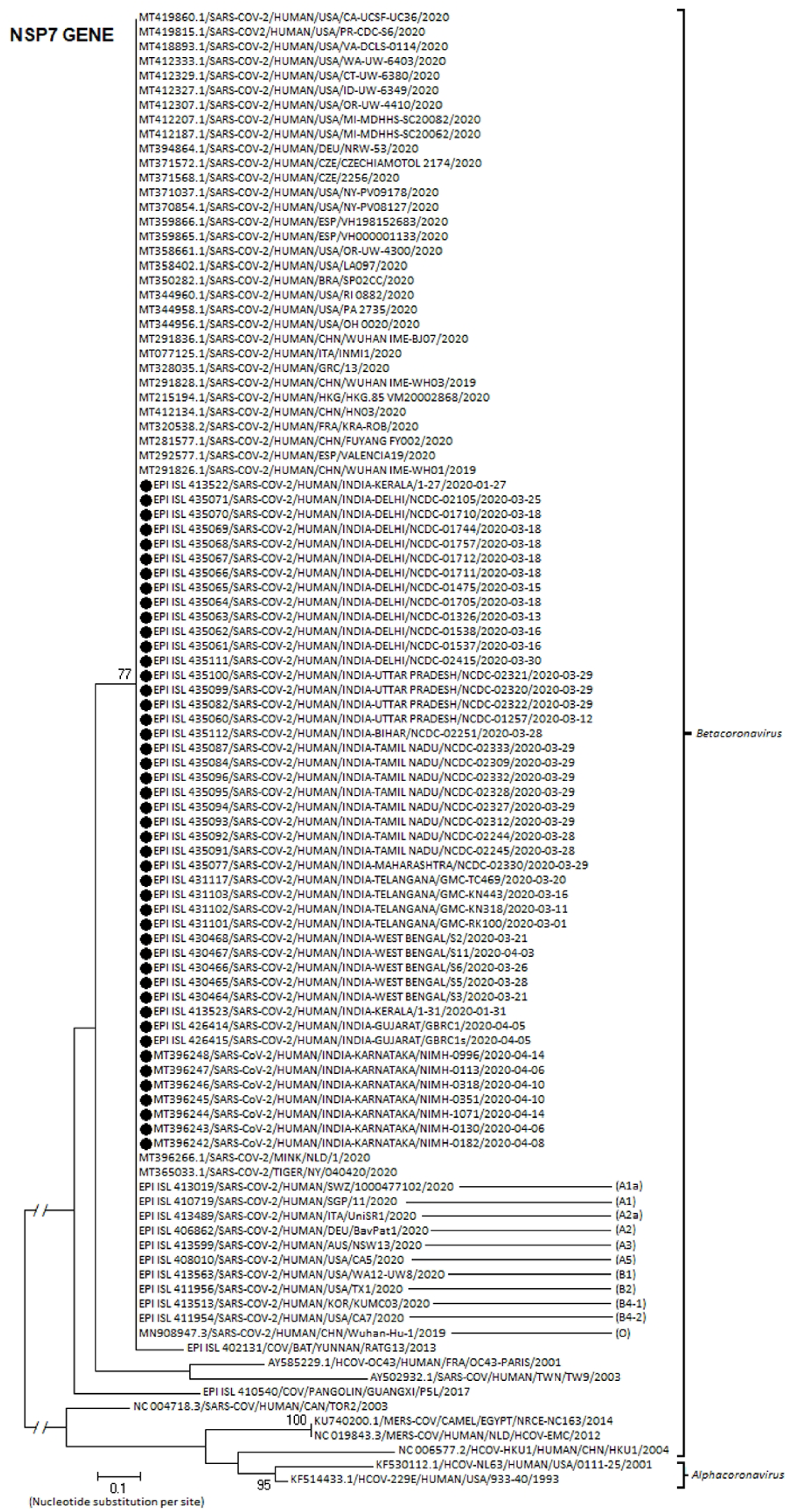
Molecular Phylogenetic analysis by Maximum Likelihood method: Phylogenetic dendrogram based on nucleotide sequences of NSP7 gene of SARS-CoV-2 strains circulating in India during early 2020, with other known strains of respective genotype. The representative Indian strains have been marked with a solid circle (●). Scale-bar was set at 0.1 nucleotide substitutions per site. Bootstrap values of less than 70% are not shown. The best fit model which was used for constructing the phylogenetic dendrogram was Tamura-3 model with Gamma distribution having Invariant sites (T92+G+I).

**Figure 1(I):**
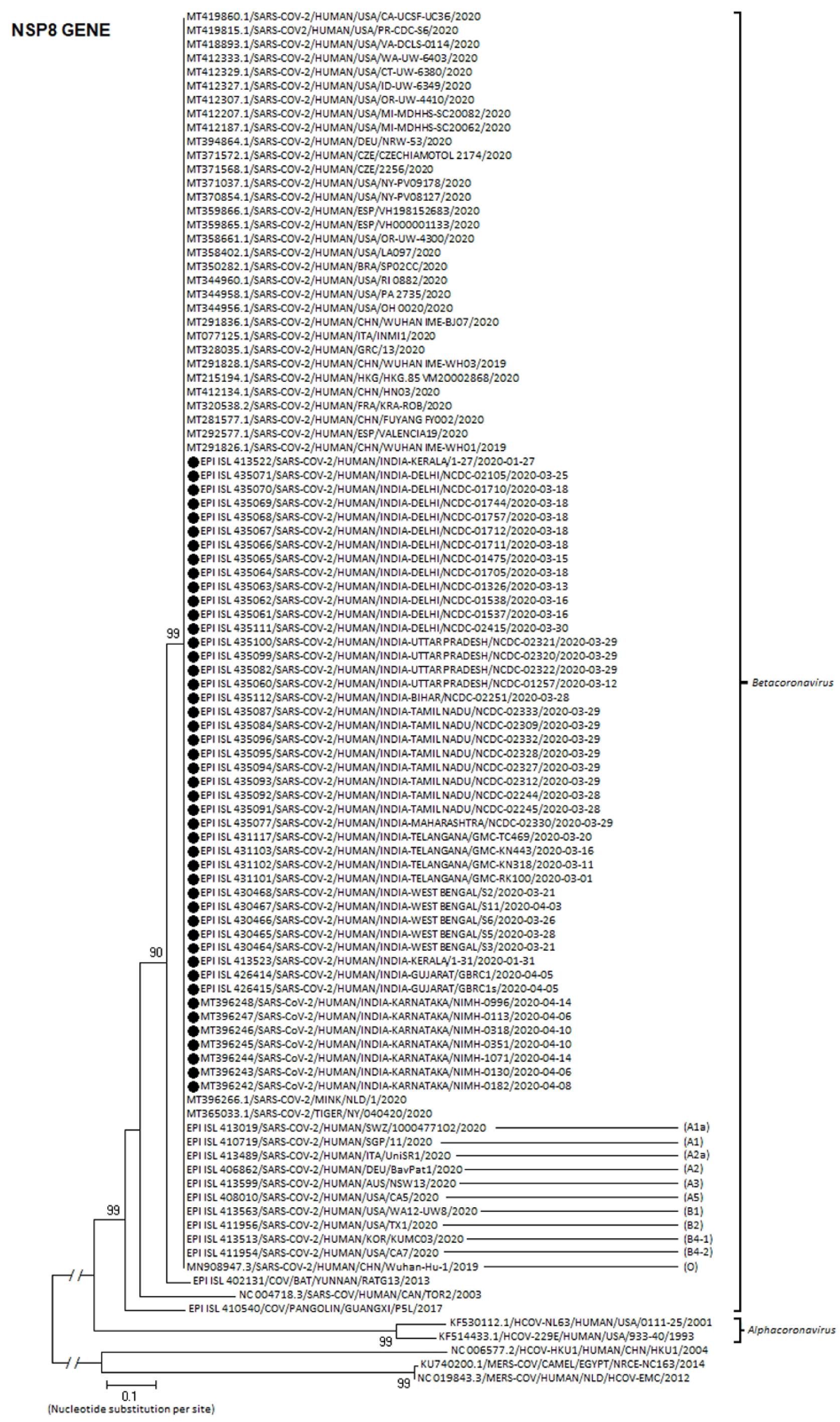
Molecular Phylogenetic analysis by Maximum Likelihood method: Phylogenetic dendrogram based on nucleotide sequences of NSP8 gene of SARS-CoV-2 strains circulating in India during early 2020, with other known strains of respective genotype. The representative Indian strains have been marked with a solid circle (●). Scale-bar was set at 0.1 nucleotide substitutions per site. Bootstrap values of less than 70% are not shown. The best fit model which was used for constructing the phylogenetic dendrogram was General Time Reversible model with Gamma distribution having Invariant sites (GTR+G+I).

### 3.2. L and S type of SARS-CoV-2

Single Nucleotide Polymorphisms (SNPs) at positions 8,782 (NSP4 gene) and 28,144 (ORF 8) showed complete linkage among the Indian isolates under study. At these two sites, 44 strains showed a “CT” haplotype (designated as “L” type as T28,144 falls inLeucine codon); while only 2 strains (one from Kerala, EPI_ISL_413523 and other from Delhi, EPI_ISL_435111) revealed a “TC” haplotype (called as “S” type as C28,144 falls in the codon for Serine) (Table 5).

**Table 1:** Single nucleotide polymorphisms associated with nonsynonymous mutations in S protein in Indian isolates during early 2020.

|  |  | S glycoprotein<br>(21563-25384 nts / 1273 amino acid) |  |  |  |  |  |
| --- | --- | --- | --- | --- | --- | --- | --- |
|  |  | Mutation 1 |  | Mutation 2 |  | Mutation 3 |  |
| Sequence accession number |  | Nucleotide position | Amino acid position | Nucleotide position | Amino acid position | Nucleotide position | Amino acid position |
|  |  | 22374<br>A>G | 271<br>Q>R | 23403<br>A>G | 614<br>D>G | 24933<br>G>T | 1124<br>G>V |
| Reference sequence | MN908947.3/SARS-COV-2 Wuhan-Hu-1 | A | Q | A | D | G | G |
| Delhi | EPI_ISL_435071 | A | Q | G | G | G | G |
|  | EPI_ISL_435070 | A | Q | G | G | G | G |
|  | EPI_ISL_435069 | A | Q | G | G | G | G |
|  | EPI_ISL_435068 | A | Q | G | G | G | G |
|  | EPI_ISL_435067 | A | Q | G | G | G | G |
|  | EPI_ISL_435066 | A | Q | G | G | G | G |
|  | EPI_ISL_435065 | A | Q | G | G | G | G |
|  | EPI_ISL_435064 | A | Q | G | G | G | G |
|  | EPI_ISL_435063 | A | Q | G | G | G | G |
|  | EPI_ISL_435062 | A | Q | G | G | G | G |
|  | EPI_ISL_435061 | A | Q | G | G | G | G |
|  | EPI_ISL_435111 | A | Q | A | D | G | G |
| Tamil Nadu | EPI_ISL_435087 | A | Q | A | D | G | G |
|  | EPI_ISL_435084 | A | Q | A | D | G | G |
|  | EPI_ISL_435096 | A | Q | A | D | G | G |
|  | EPI_ISL_435095 | A | Q | A | D | G | G |
|  | EPI_ISL_435094 | A | Q | A | D | G | G |
|  | EPI_ISL_435093 | A | Q | A | D | G | G |
|  | EPI_ISL_435092 | A | Q | A | D | G | G |
|  | EPI_ISL_435091 | A | Q | A | D | G | G |
| Karnataka | MT396248 | A | Q | A | D | G | G |
|  | MT396247 | A | Q | G | G | G | G |
|  | MT396246 | A | Q | A | D | G | G |
|  | MT396245 | A | Q | A | D | G | G |
|  | MT396244 | A | Q | A | D | G | G |
|  | MT396243 | A | Q | G | G | G | G |
|  | MT396242 | A | Q | G | G | G | G |
| West Bengal | EPI_ISL_430468 | A | Q | G | G | T | V |
|  | EPI_ISL_430467 | A | Q | G | G | G | G |
|  | EPI_ISL_430466 | A | Q | G | G | G | G |
|  | EPI_ISL_430465 | A | Q | G | G | G | G |
|  | EPI_ISL_430464 | A | Q | G | G | T | V |
| Uttar Pradesh | EPI_ISL_435100 | A | Q | A | D | G | G |
|  | EPI_ISL_435099 | A | Q | A | D | G | G |
|  | EPI_ISL_435082 | A | Q | A | D | G | G |
|  | EPI_ISL_435060 | A | Q | G | G | G | G |
| Telangana | EPI_ISL_431117 | A | Q | G | G | G | G |
|  | EPI_ISL_431103 | A | Q | A | D | G | G |
|  | EPI_ISL_431102 | A | Q | G | G | G | G |
|  | EPI_ISL_431101 | A | Q | A | D | G | G |
| Gujarat | EPI_ISL_426414 | G | R | G | G | G | G |
|  | EPI_ISL_426415 | G | R | G | G | G | G |
| Kerala | EPI_ISL_413522 | A | Q | A | D | G | G |
|  | EPI_ISL_413523 | A | Q | A | D | G | G |
| Bihar | EPI_ISL_435112 | A | Q | A | D | G | G |
| Maharashtra | EPI_ISL_435077 | A | Q | A | D | G | G |

**Table 2:**
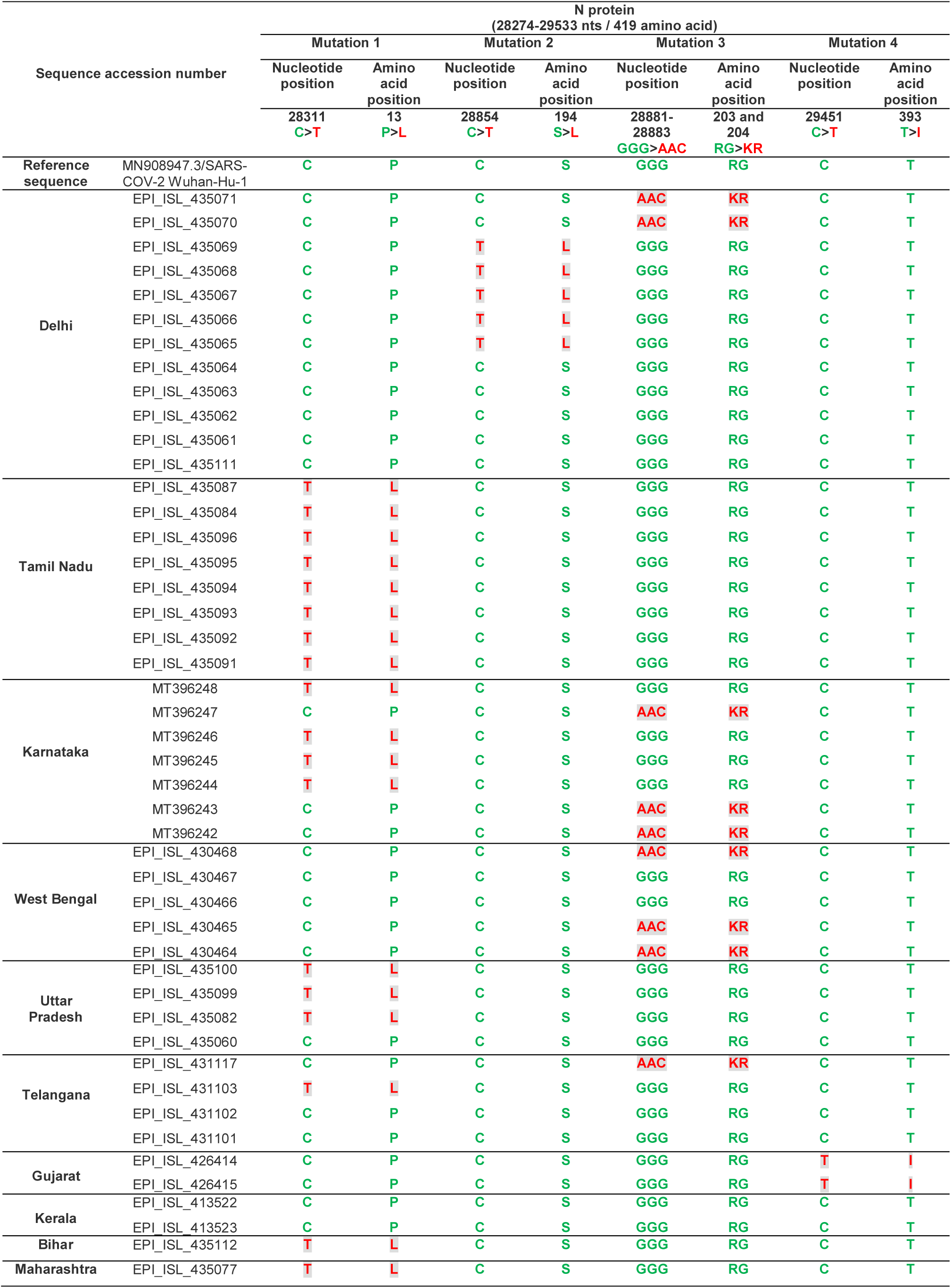
Single nucleotide polymorphisms associated with nonsynonymous mutations in N protein in Indian isolates during early 2020.

**Table 3:** Single nucleotide polymorphisms associated with nonsynonymous mutation(s) in NSP6 and RdRP proteins in Indian isolates during early 2020.

| Sequence accession number |  | NSP6 protein<br>(10973-11842 nts / 290 amino acid) |  | RdRP protein<br>(13442-16236 nts / 932 amino acid) |  |  |  |
| --- | --- | --- | --- | --- | --- | --- | --- |
|  |  | Mutation 1 |  | Mutation 1 |  | Mutation 2 |  |
|  |  | Nucleotide position | Amino acid position | Nucleotide position | Amino acid position | Nucleotide position | Amino acid position |
|  |  | 11083<br>G>T | 37<br>L>F | 13730<br>C>T | 97<br>A>V | 14408<br>C>T | 323<br>P>L |
| Reference sequence | MN908947.3/SARS-COV-2 Wuhan-Hu-1 | G | L | C | A | C | P |
| Delhi | EPI_ISL_435071 | G | L | C | A | T | L |
|  | EPI_ISL_435070 | G | L | C | A | T | L |
|  | EPI_ISL_435069 | G | L | C | A | T | L |
|  | EPI_ISL_435068 | G | L | C | A | T | L |
|  | EPI_ISL_435067 | G | L | C | A | T | L |
|  | EPI_ISL_435066 | G | L | C | A | T | L |
|  | EPI_ISL_435065 | G | L | C | A | T | L |
|  | EPI_ISL_435064 | G | L | C | A | T | L |
|  | EPI_ISL_435063 | G | L | C | A | T | L |
|  | EPI_ISL_435062 | G | L | C | A | T | L |
|  | EPI_ISL_435061 | G | L | C | A | T | L |
|  | EPI_ISL_435111 | G | L | C | A | C | P |
| Tamil Nadu | EPI_ISL_435087 | T | F | T | V | C | P |
|  | EPI_ISL_435084 | T | F | T | V | C | P |
|  | EPI_ISL_435096 | T | F | T | V | C | P |
|  | EPI_ISL_435095 | T | F | T | V | C | P |
|  | EPI_ISL_435094 | T | F | T | V | C | P |
|  | EPI_ISL_435093 | T | F | T | V | C | P |
|  | EPI_ISL_435092 | T | F | T | V | C | P |
|  | EPI_ISL_435091 | T | F | T | V | C | P |
| Karnataka | MT396248 | G | L | Not taken for analysis due to poor sequence reads |  |  |  |
|  | MT396247 | G | L |  |  |  |  |
|  | MT396246 | T | F |  |  |  |  |
|  | MT396245 | G | L |  |  |  |  |
|  | MT396244 | G | L |  |  |  |  |
|  | MT396243 | G | L |  |  |  |  |
|  | MT396242 | G | L |  |  |  |  |
| West Bengal | EPI_ISL_430468 | G | L | C | A | T | L |
|  | EPI_ISL_430467 | G | L | T | V | T | L |
|  | EPI_ISL_430466 | G | L | C | A | T | L |
|  | EPI_ISL_430465 | G | L | C | A | T | L |
|  | EPI_ISL_430464 | G | L | C | A | T | L |
| Uttar Pradesh | EPI_ISL_435100 | T | F | T | V | C | P |
|  | EPI_ISL_435099 | T | F | T | V | C | P |
|  | EPI_ISL_435082 | T | F | T | V | C | P |
|  | EPI_ISL_435060 | G | L | C | A | T | L |
| Telangana | EPI_ISL_431117 | G | L | C | A | T | L |
|  | EPI_ISL_431103 | T | F | T | V | C | P |
|  | EPI_ISL_431102 | G | L | C | A | T | L |
|  | EPI_ISL_431101 | G | L | C | A | C | P |
| Gujarat | EPI_ISL_426414 | G | L | C | A | T | L |
|  | EPI_ISL_426415 | G | L | C | A | T | L |
| Kerala | EPI_ISL_413522 | G | L | C | A | C | P |
|  | EPI_ISL_413523 | G | L | C | A | C | P |
| Bihar | EPI_ISL_435112 | T | F | T | V | C | P |
| Maharashtra | EPI_ISL_435077 | T | F | T | V | C | P |

**Table 4:**
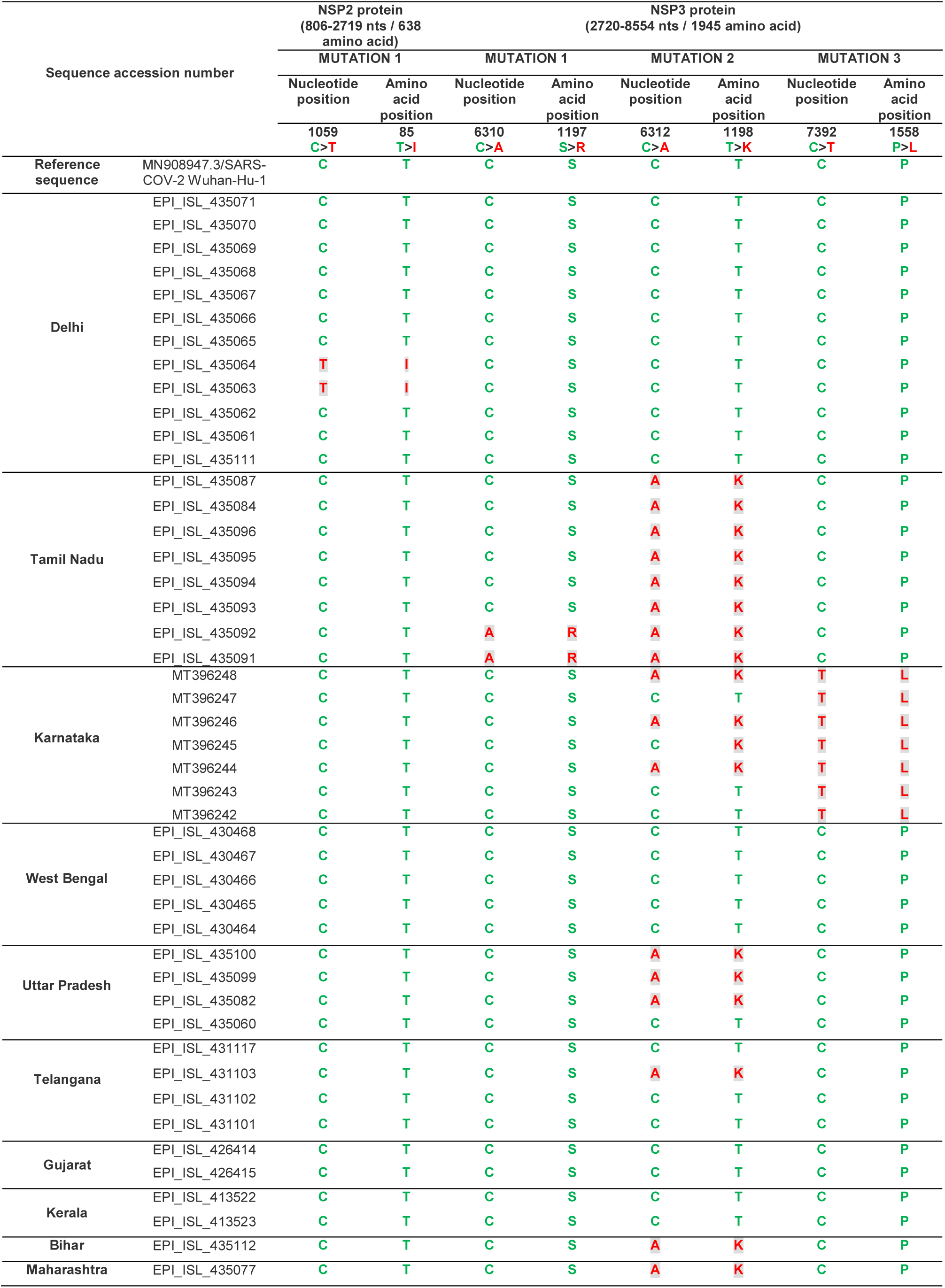
Single nucleotide polymorphisms associated with nonsynonymous mutation(s) in NSP2 and NSP3 proteins in Indian isolates during early 2020.

**Table 5:**
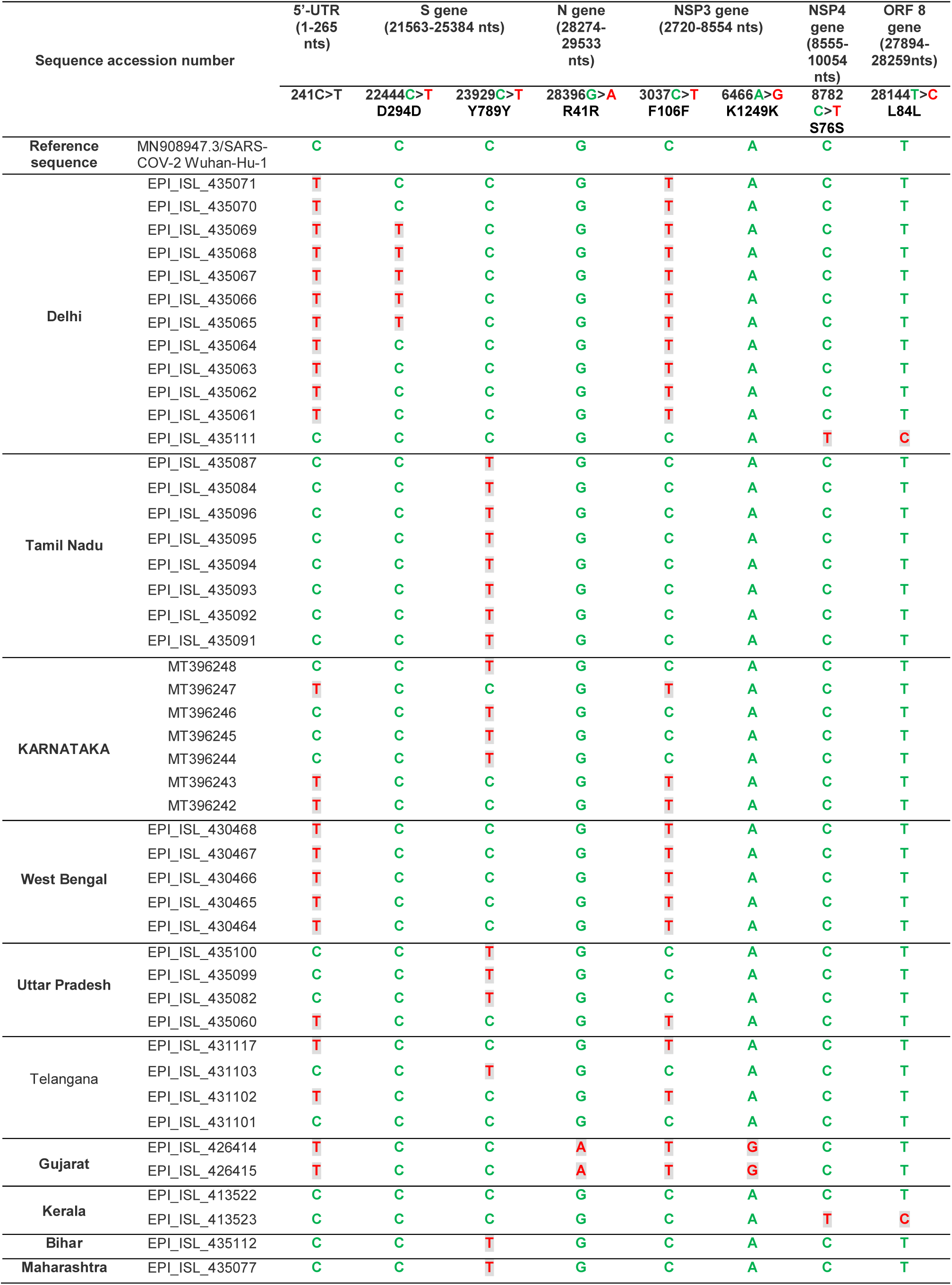
Single nucleotide polymorphisms associated with 5’-UTR or synonymous mutation(s) in S, N, NSP3, NSP4 and ORF8 proteins in Indian isolates during early 2020.

### 3.3. Analyses of Synonymous and Non-Synonymous Mutations

#### 3.3.1. The common mutations in SARS-CoV-2 Indian isolates

In order to explore the mutationsamong the 46 SARS-CoV-2 strains, we performed intricate sequence analyses both at the genome level and corresponding amino acid level in different proteins, especially Spike glycoprotein (S), Nucleocapsid protein (N), Non-structural protein 2 (NSP2), NSP3, NSP4, NSP6, NSP7, NSP8 and RNA-dependent RNA polymerase (RdRP/NSP12) with reference to the prototype SARS-CoV-2 strain (MN908947.3/SARS-COV-2/HUMAN/CHN/Wuhan-Hu-1/2019). 4 out of 46 samples (8.69%) lacked any significant mutations: one collected from a patient with unknown travel history (EPI_ISL_435111), one with travel history to Dubai (EPI_ISL_431101) and two patients with travel history to China (EPI_ISL_413522 and EPI_ISL_413523). Mutational analysis of remaining 42 strains revealed circulation of two predominant “groups”, namely the “major group” and the “minor group”, in India. The “major group” which comprised of 24 isolates (24/46, 52.17%), revealed four co-existing SNPs; 241C>T in the 5-UTR (Table 5), 3037C>T (F106F) in the NSP3 gene (Table 5), 14408C>T (P323L) in NSP12 gene (Table 3, Mutation 2) and 23403A>G (D614G) in the S gene (Table 1, Mutation 2). This “major group” of SARS-CoV-2 was predominantly found to circulate in Delhi, West Bengal, Telangana and Gujarat. Rest 18 samples (18/46, 39.13%) were seen to have three co-existing mutations; 23929C>T (Y789Y) in the S gene (Table 5), 28311C>T (P13L) in the N gene (Table 2, Mutation 1) and 6312C>A (T1198K) in the NSP3 gene (Table 4, Mutation 2). Out of these 18 isolates, 14 samples (14/46, 30.43%) that represent the “minor group”had two additional co-existing mutations; 11083G>T (L37F) in the NSP6 gene (Table 3, Mutation 1, marked with *) and 13730C>T (A97V) in the NSP12/RdRP gene (Table 3, Mutation 1, marked with ¶). Needless to say, the five co-existing mutations of the “minor group” and the four co-existing mutations of the “major group” were not overlapping among the same SARS-CoV-2 isolate. The “minor group” of SARS-CoV-2 predominated in Tamil Nadu, Karnataka, Uttar Pradesh, Bihar and Maharashtra.

#### 3.2.2. The unique mutations in SARS-CoV-2 Indian isolates

In addition to 23403A>G (D614G), three uncommon mutations 23374A>G (Q271R) (Table 1, Mutation 1), 24933G>T (G1124V) (Table 1, Mutation 3) and 22444C>T (D294D) (Table 5) were also observed in the S gene of “major group”. 16 out of the 24 isolates revealed three novel mutations 28854C>T (S194L) (5 samples), 28881-28883GGG>AAC (R203K and G204R) (9 samples) and 29451C>T (T393I) (2 samples) in the N gene (Table 2, Mutation 2, 3 and 4 respectively), whereas two isolates of the “minor group”had 28396G>A (R41R) change in the N gene (Table 5). Intriguingly, 28854C>T (S194L) in N gene was found to co-evolve with 22444C>T (D294D) mutation inthe S gene. We also observed 1059T>A (T85I) change within the NSP2 (Table 4) and, 6310C>A (S1197R), 7392C>T (P1558L) (Table 4) and 6466A>G (K1249K) (Table 5)change in the NSP3 gene. No mutations were found within the NSP7 and NSP8 genes.

#### 3.3.3. Effect of missense mutation A97V on the secondary structure of NSP12/RdRP

Being the crucial enzyme for viral RNA replication and maintaining the genomic fidelity, any significant change in RdRP structure could affect its functions, thereby increasing the rate of mutagenesis in the genome. We have identified two missense mutations in the RdRP protein; P323L associated with the “major group” isolates and A97V associated with the “minor group” isolates. The effect of P323L on the secondary structure of RdRP has already been described [8]. Therefore, we analyzed the effect of novel mutation A97V on the secondary structure of RdRP by using CFSSP server. A97V mutation resulted in substitution of α-helixes at positions 94, 95 and 96 with the β-sheets in the RdRP secondary structure which might alter its tertiary conformation, resulting in significant functional implications (Fig.2).

**Figure 2:**
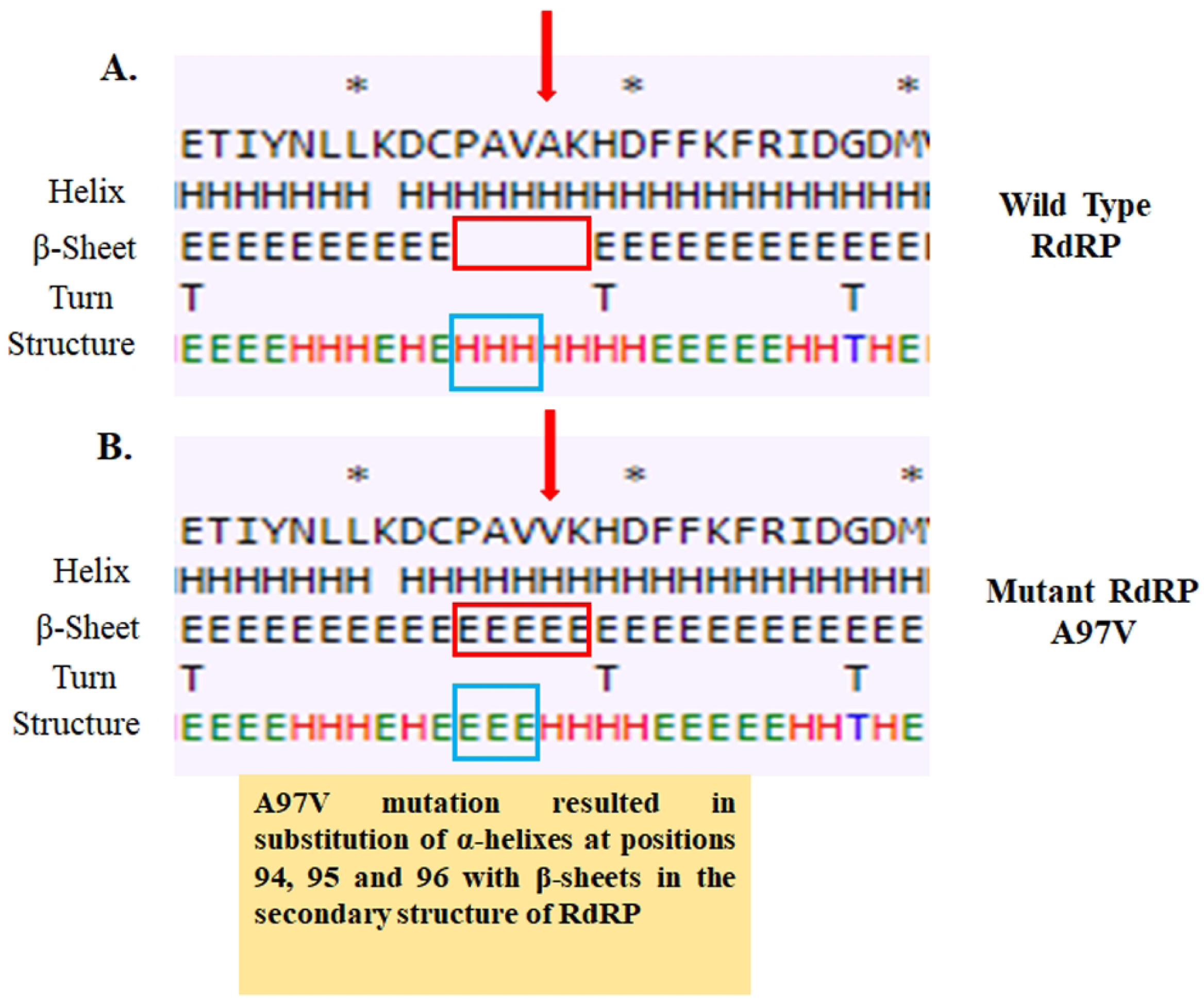
Effect of A97V mutation on the secondary structure of RdRP/ NSP12 protein. (A) Secondary structure of RdRP around 97^th^ A (Alanine) residue of Wuhan isolate of SARS-CoV-2. (B) Secondary structure of RdRP around 97th V (Valine) residue of Indian isolate of SARS-CoV-2.

## 4. Discussion

Molecular and genetic characterization of SARS-CoV-2 pandemic strains worldwide has been studied by several scientific groups based on whole-genome sequencing [9,10]. Through this comprehensive analysis we aimed to closely analyze the ancestry, evolutionary dynamics, accumulation of rapid mutations and cross-genetic translation among the emerging SARS-CoV-2 strains across India. As depicted through the different phylogenetic dendrograms in our study, a monophyletic clade of all SARS-CoV-2 strains was seen with the prototype strain (Wuhan IME-WH01/2019) clustering along with the Indian isolates. Rapid accumulation of several point mutations across the genome of SARS-CoV-2 since its origin is a prime driving force behind the evolution of different monophyletic clades. Clustering of all the India isolates with other SARS-CoV-2 strains reported worldwide (99.8-100% nucleotide sequence identity) suggests forward introduction of this virus in India from several countries. Clustering of the prototype strain from Wuhan underscores the fact that China might have served as the origin of this zoonotic virus, which has been eventually transmitted to worldwide [11-13].

The origin of the SARS-CoV-2 is still a debatable issue but identification of its intermediate host is much needed to prevent further dissemination and interspecies transmission in near future. Hence, we initiated this study as one of the first in India to decipher the gene-wise phylogenetics of SARS-CoV-2 strains circulating in this endemic setting. The results depicted genome type clusters of the 46 Indian isolates, for the structural genes S and N and the non-structural gene RdRP/NSP12. Clustering of the study isolates with different clade-specific strains for different gene establishes the development of genome type clustering. Though variations in DNA homology exists with respect to each gene, nevertheless a very recent bifurcation of these Indian strains from the bat and Malayan-pangolin derived SARS-like coronaviruses is supposed to have occurred, with a subsequent zoonotic transmission to humans, as depicted through all the nine dendrograms. However, SARS-CoV-2 strains were distant to MERS-CoV and other human coronaviruses. This conclusion goes at par with other phylogenetic studies establishing bat and pangolins as the proximal origin of SARS-CoV-2 [14-16].

Our study highlighted low sequence similarity of spike gene with some bat derived strains like bat-SL-CoVZC45 and bat-SL-CoVZXC21, while maximum homology was noticed with bat SARS-like Coronavirus (SARSr-CoV/RaTG13). In contrast to a study by Zhang et.al, the pangolin-derived coronavirus revealed very low sequence homology to all the Indian SARS-CoV-2 strains than the bat RaTG13 [2]. The RNA Binding Domain (RBD) within the S1 subunit of Spike gene of all Indian SARS-CoV-2 and Pangolin-derived strains were found to be evolutionarily conserved and phylogenetically much closer than bat RaTG13, underscoring familiar mode of pathogenesis between the two. The Indian SARS-CoV-2 isolates too possess a polybasic cleavage site (RRAR; amino acid position 682-685) at the junction of S1 and S2 subunits of Spike protein as reported by Andersen et.al., 2020 [14]. SARS-CoV2 strains have been categorized into two major groups/types characterized by two SNPs at positions 8782 (NSP4 gene) and 28,144 (ORF8) that reveal complete linkage [17]. Among our Indian study isolates, frequency of the L-type (CT haplotype) was much higher (44/46, 95.7%) to the S-type (TC haplotype) (2/46, 4.3%), indicating the L-type to predominate over S-type in this geographical region.

Convoluted mutational analysis also revealed co-circulation of two groups of mutated SARS-CoV-2 strains in India. The “major group” of SARS-CoV-2 strains (52.2%) represents a modified A2a clade reported previously from Africa, South America, Oceania and South and West Asia comprising of strains with co-evolving mutations like 241 C>T (5’ UTR), 3037 C>T (F106F, NSP3), 14403 C>T (P323L, RdRP/ NSP12) and 23403 A>G (D614G, S glycoprotein) [18-20]. Certain strains in the “major group” displayed 22374A>G (Q271R), 24933G>T (G1124V) and 22444C>T (D294D) changes in the S gene which are unique to India. Missense mutations Q271R and G1124V in S protein reside around the N-linked glycosylation sites 282 and 1134 respectively and these might affect the protein function [21]. It was not surprising to observe the triple site mutation 28881-28883 GGG>AAC (R203K and G204R) in the N gene of 9 SARS-CoV-2 strains of the “major group”. This has previously been reported from Mexico, South America, Australia, New Zealand and few Asian countries as well [22]. The 203/204 region is part of the SR dipeptide domain of N protein (**SR**NS**SR**NSTPGS**SR**GTSPARMA) and changes in Arg^203^ to Lys^203^ and Gly^204^ to Arg^204^ resulted in the insertion of a lysine residue between serine and arginine (**SR**NS**SR**NSTPGS**SKR**TSPARMA) which might interfere with the phosphorylation at serine residue required for normal functioning of the N protein [23]. This mutation demands particular attention as reduced pathogenicity has been observed in SARS-CoV on deletion of the SR domain [24]. Mutations observed in the NSP3 gene at positions 6310 C>A (S1197R), 7392 C>T (P1558L) and 6466 A>G (K1249K) were completely unique to Indian strains. Few infrequent mutations at position 1059 T>A (T85I) in NSP2 and 8782 C>T (S76S) in NSP4 have also been reported to be prevalent in other countries [19, 22, 25].

The “minor group” of Indian SARS-CoV-2 (30.4%), comprised of strains with five co-evolving mutations: 13730C>T (A97V, RdRP/NSP12), 23929C>T (Y789Y, S), 28311C>T (P13L, N), 6312C>A (T1198K, NSP3) and 11083G>T (L37F, NSP6). All the “minor group” mutations were novel among the Indian isolates, except 11083G>T (L37F, NSP6) which was previously reported as infrequent mutation from Australia, Japan, Netherlands and some other European countries [18,26]. The L37F mutation strongly implies positive selection towards evolution of betacoronaviruses, indicating a possible origin of the “minor group” out of this positive selection and subsequent acquisition of mutations among the strains already harboring the 11083G>T change [25,26]. The interaction of NSP6 with NSP3 and NSP4 has been described to be essential for the formation of double membrane vesicles [25,26]. Hence, it is very interesting to note the presence of a co-existing mutation 6312 C>A (T1198K) in NSP3 of the “minor group” strains, though the significance of this co-existence (L37F and T1198K) in context toNSP6-NSP3 interaction can only be confirmed through association studies. The fidelity of RdRP is challenged due to the presence of 13730 C>T (A97V) change, which was predicted to have significant effect on the secondary structure of RdRP. A97V was found to be located in the nidovirus RdRP-associated nucleotidyltransferase (NiRAN) domain whose function remains unknown [27]. The P13L mutation is located in the intrinsically disordered region of the N protein and might affect RND-binding activity of N-terminal domain (NTD) and C-terminal domain (CTD) of the N protein [28, 29].

Any significant mutation in the RdRP/NSP12 protein might alter replication machinery, thereby compromising the fidelity of viral RNA replication and subsequent accumulation of plausiblenovel mutations. The missense mutation 14408C>T (P323L) in RdRP was first observed in Italy (Lombardy) in February, 2020. The study by Pachetti et al. has shown emergence of 3037C>T (F106F, NSP3 protein), 23403A>G (D614G) and 28881-28883GGG>AAC (R203K and G204R, N protein) in SARS-CoV-2 having 14408C>T (P323L) in RdRP gene February, 2020 in Europe and North America, suggesting a probable association of 14408C>T (P323L) and the emergence of other new mutations [30]. Therefore, we can assume that two the mutations: 14408C>T (P323L) and 13730C>T (A97V), which were found to have significant influence on the secondary structure of RdRP, could play key roles in the simultaneous establishment of “two groups” of SARS-CoV-2 with characteristic “co-evolving mutations” in India. However, this needs to be validated experimentally. Owing to lack of patient metadata, we could not determine whether these two distinct groups of strains varied in their virulence and pathogenicity.

This study adds up few initial observations on the available Indian SARS-CoV-2 sequences like ancestry, evolutionary dynamics, clade specific genetic variations as well as development of unique co-evolving mutations and their possible effect on protein secondary structure. The ongoing deadly pandemic demands an urgent need to record complete patient metadata along with full genome sequences of the SARS-CoV-2 strain in varied epidemiological settings for better understanding of pathogenesis of disease and its correlation with accumulated mutation in viral genome.

## Acknowledgement

This study was supported by intramural grant from the Indian Council of Medical Research, New Delhi, India.

The authors acknowledge the hard work and dedication of scientists performing next generation genome sequencing and submitting them in public data bases for the benefit of scientific community.

Rakesh Sarkar and Mahadeb Lo were supported by fellowships from University Grants Commission and Council of Scientific and Industrial Research, India, respectively.

